# RANbodies: reporter-nanobody fusions as versatile, small, sensitive immunohistochemical reagents

**DOI:** 10.1101/238162

**Authors:** Masahito Yamagata, Joshua R. Sanes

## Abstract

Sensitive and specific antibodies are essential for detecting molecules in cells and tissues. However, currently used polyclonal and monoclonal antibodies are often less sensitive than desired, difficult to produce, and available in limited quantities. A promising recent approach to circumvent these limitations is to employ chemically-defined antigen-combining sites called nanobodies, derived from single chain camelid antibodies. Here, we used nanobodies to prepare sensitive unimolecular detection reagents by genetically fusing cDNAs encoding nanobodies to enzymatic or antigenic reporters. We call these fusions between a **r**eporter **a**nd a **n**anobody RANbodies. They can be used to localize epitopes and to amplify signals from fluorescent proteins. They be generated and purified simply and in unlimited amounts, and can be preserved safely and inexpensively in the form of DNA or digital sequence.

## INTRODUCTION

Sensitive, specific and reproducible localization of molecules in cells and tissues is indispensable in many areas of biological inquiry. The most commonly used methods are immunohistochemical. The introduction of the immunofluorescent technique, in which antibodies are conjugated to a fluorophore (1) had a profound influence, but suffered from limited sensitivity and specificity, the need to laboriously generate conjugates of each antibody preparation, and the potential loss of activity upon conjugation. These limitations were addressed by refinements of the technology, including affinity purification of monospecific antibodies from sera to improve specificity, use of enzymatic labels such as horseradish peroxidase to improve sensitivity, and use of second antibodies (indirect immunofluorescence) to avoid the need for chemical modification of the primary antibody (2, 3).

Nonetheless, difficulties remained. Polyclonal antibodies from immunized animals are often poorly defined, available in limited amounts and vary from bleed to bleed (4, 5). Some but not all of these problems were addressed by the introduction of monoclonal antibodies generated from hybridomas (6); they are monospecific, molecularly defined, and can be produced in unlimited amounts. However, they are laborious to generate. Moreover, storage of hybridomas is costly and, even at low temperatures, impermanent.

A subsequent step toward routine production of specific and sensitive immunoreagents was the development of methods for selection and generation of recombinant antigen-binding antibody fragments. The first of these were variable regions from conventional antibodies, cloned by PCR and then evolved, and selected by methods such as phage display (7, 8). More recently, single chain antibodies from camelids (9) or selachians (10), have been used for research, diagnostic, and therapeutic purposes (11). Importantly, the high-affinity of the antigen recognition site in these single-chain molecules is retained in fragments called nanobodies that comprise only the ~100 amino-acid long variable domain enabled (12, 13). Nanobodies are thus easy to clone and derivatize by coupling to reporters or dyes (14–17). Moreover, they can be stored with high stability and low cost as cDNAs, or regenerated at relatively low cost from digitally stored sequence.

Here, we report an immunohistochemical platform based on nanobodies. We fuse the nanobody to a reporter and append an epitope tag that enables detection of the protein independent of its bioactivity, as well as one-step affinity purification from culture medium. Nanobody sequences can be synthesized and cloned into a destination vector containing all other elements. We call these detection RANbodies for fusions between a **r**eporter **a**nd a **n**anobody. Here, we describe methods for generating RANbodies using each of four reporters: a variant of horseradish peroxidase, two highly antigenic proteins (“spaghetti monsters”, ref. 18) and the Fc fragment of an avian antibody. We document the versatility of these reagents for detection of antigens in cultured cells and tissues, with an emphasis on amplifying weak signals from multiple fluorescent proteins (XFPs). We believe that the simplicity of the method and specificity of the reagents will make them widely useful.

## RESULTS

### RANbody platform

RANbody probes contain the following 6 elements: (a) an N-terminal mammalian signal peptide from human immunoglobulin kappa chain to enable secretion of the protein from cultured cells; (b) a hemagglutinin (HA) epitope tag to enable immunochemical detection of the protein independent of its binding and enzymatic activities; (c) a camelid nanobody; (d) a short linker; (e) a reporter; and (f) a histidine (His) epitope tag to enable one-step affinity purification of the protein from culture medium (Fig. 1*A, B*).

**Fig. 1.**
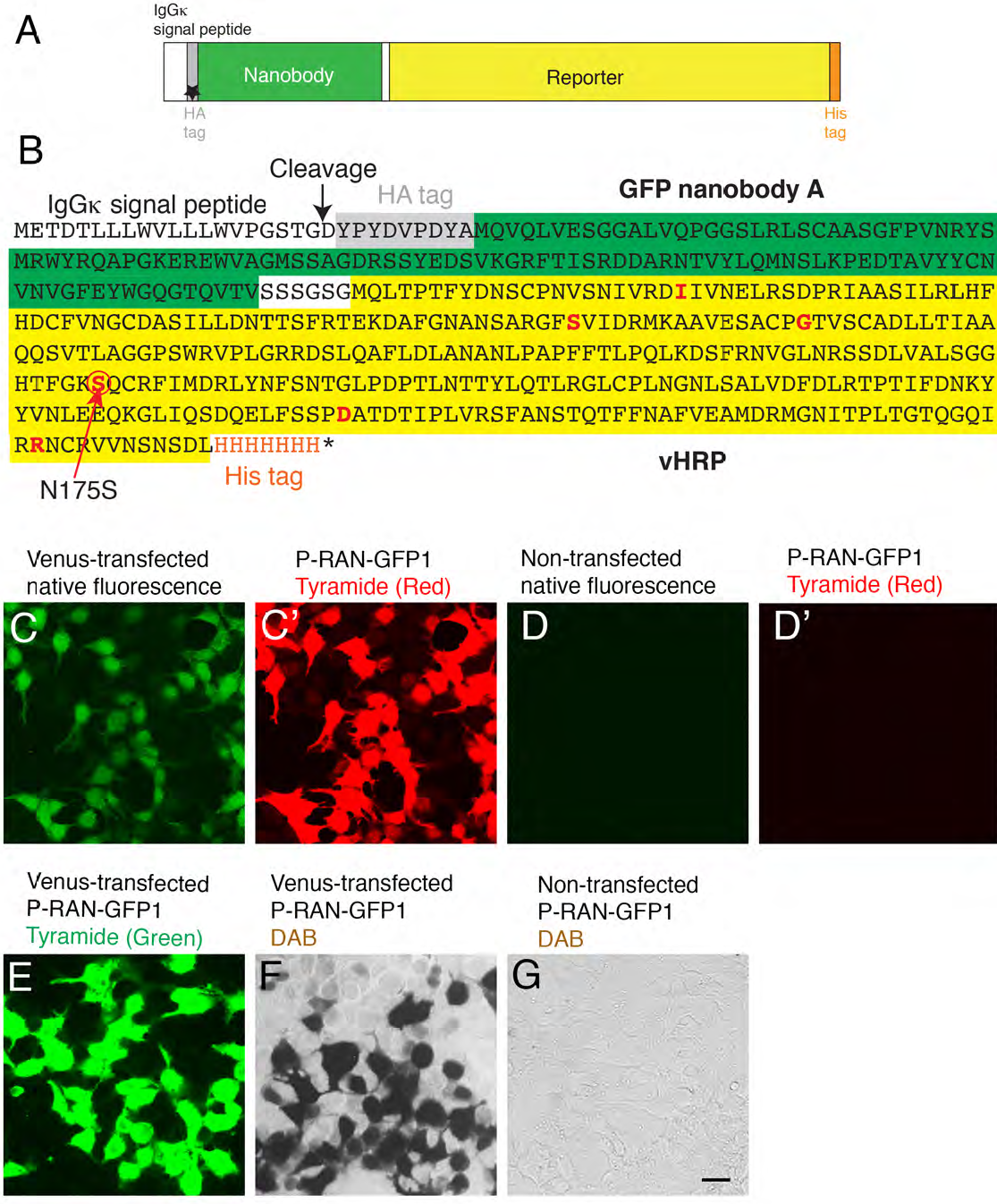
Structure and use of RANbodies. (*A*) RANbody structure. Elements, in order, are (a) a mammalian signal sequence, (b) hemagglutinin (HA) epitope tag, (c) nanobody, (d) spacer, (e) reporter, (f) stretch of small amino acids, and (g) polyhistidine (His) epitope tag. (*B*) Sequence of P-RAN-GFP1 The reporter is an enhanced variant of HRP (vHRP). Amino acids in red show mutations that enhance activity compared to native HRP; of these N175S has the greatest effect. (*C,C*’) RANbody staining. 293T cells transfected with a YFP variant (Venus) were incubated with GFP-RAN-P1 and stained with tyramide-Cy3) (*C*, native fluorescence; *C*’, Cy3 fluorescence). (*D,D*’) Untransfected cells stained as in c. (*E*) Transfected cells stained as in c but with tyramide-FITC. (*F*) Transfected cells stained as in c but with a chromogenic substrate, diaminobenzidine (DAB). (*G*) Untransfected cells stained as in *F*. Bar, 10 *μ*m.

For facile construction of RANbodies, we used the Gibson assembly method (19), in which multiple overlapping DNA molecules can be assembled in a single step. The nanobody fragments, which are ~400 bp long, were synthesized commercially and other components were generated by PCR from readily available plasmids. The resulting vectors were transfected into the mammalian cell line, 293T, and RANbodies were collected from the medium. Diluted culture medium was adequate for staining in most cases. However, by using the His tag at the carboxy terminus, we were able to concentrate the activity by more than two orders of magnitude in a single step on a commercially available affinity resin, as judged by peroxidase activity. The purified recombinant proteins (~450 amino acids) migrated as a single ~60-65 kDa band on SDS-polyacrylamide gels (Fig. S1). The diffuse nature of the band likely reflects heterogeneity in glycosylation.

### Nomenclature

In describing the construction and use of RANbodies, we use a simple nomenclature in which the term “RAN” is preceded by a one letter abbreviation denoting the reporter and followed by the antigen to which the nanobody is directed. Reporters described here are horseradish peroxidase (P), HA-tagged spaghetti monster (H), Myc-tagged spaghetti monster (M) and chicken IgY-Fc region (Y). Thus, a RANbody incorporating a nanobody directed at GFP and horseradish peroxidase is denoted P-RAN-GFP. If there are multiple RANbodies with this design, they can be distinguished by number, for example P-RAN-GFP1, P-RAN-GFP2, etc.

### Optimization of HRP as reporter

The plant derived horseradish peroxidase (HRP) is a highly sensitive and commonly used reporter (20). We therefore initially used HRP as a reporter. To this end, we compared three HRP variants: The first, erHRP, was a “humanized” version of the horseradish protein, with no changes to the amino sequence, but with its nucleotide sequence optimized to improve translation efficiency in mammalian cells. An endoplasmic reticulum retention signal was appended to the carboxy terminus to allow folding and glycosylation of the protein, which does not occur in the cytoplasm (21). The second variant, sHRP, bore a single mutation, N175S, which enhances enzyme activity and protein stability (22, 23). The third variant, vHRP, bears additional five point mutations, which were selected to improve the stability of a reconstituted “split HRP” (24) but had not been studied as a single enzyme. All three versions were fused to an HA tag, so that protein concentration could be compared. Of the three, vHRP was most active, whether measured histochemically (Fig. S2*A*) or by enzyme assay in solution (Fig. S2 *B* and *C*). This version was therefore used in to generate RANbodies of the P series (Fig. 1*A*, 1*B* and Table S1).

### P-RANbody: Detection of GFP with HRP-containing RANbody

We first generated and purified a RANbody bearing a GFP nanobody described by Kubala et al., (25). Using the nomenclature described above, we refer to this reagent as P-RAN-GFP1. We used P-RAN-GFP1 to stain 293T cells transfected with Venus, an enhanced yellow fluorescent protein modified from *Aequorea victoria* GFP (26). Cells were fixed, permeabilized, incubated with RANbody overnight, then rinsed and incubated with an HRP substrate. Venus was readily detected using a fluorogenic tyramide substrate with red (Cy3) or green (FITC) fluorescence (Fig. 1*C-E*) or a chromogenic HRP substrate, 3, 3'-diaminobenzidine (DAB), which generates visible brown precipitates (Fig. 1*F*). Untransfected cells were unstained (Fig. 1*G*).

### Comparison of P-RANbody and antibody detection

We compared the sensitivity of P-RAN-GFP1 to intrinsic GFP (Venus) fluorescence and conventional indirect immunofluorescence in two ways. First, we imaged fields of 293T-transfected cells that had been stained with P-RAN-GFP1 and a red tyramide substrate. Some transfected cells were readily identified with the RANbody even though intrinsic fluorescence was barely detectable (Fig. S3*A*). Second, we stained parallel samples with monoclonal or polyclonal antbodies to GFP, followed by an appropriate second antibody, or with P-RAN-GFP1, then imaged them using confocal optics with equal gain (Fig. S3*B*). At the lowest gain, only RANbody-stained cells were visible. At intermediate gain, both RANbody and antibody-stained cells were visible. Only with the highest gain, when RANbody-stained samples were highly saturated, was intrinsic fluorescence detectable. Comparison of images acquired at the same gain indicated that RANbody increased signals ≥10-fold over antibody staining and ≥100-fold over intrinsic fluorescence.

### Detection of cellular antigens

To ask whether RANbodies could also be used to localize endogenous proteins, we generated reagents that recognized the histone H2A/H2B heterodimer and the active-binding protein, gelsolin, using published nanobody sequences (27, 28) (Table S1). Histones are confined to nuclei, and gelsolin is cytoplasmic. As expected, P-RAN-H2A2B stained nuclei in 293T cells and P-RAN-Gelsolin stained the cytoplasm using either fluorescent (Fig. 2*A-C*) or colorimetric (Fig. 2*E*) detection.

**Fig. 2.**
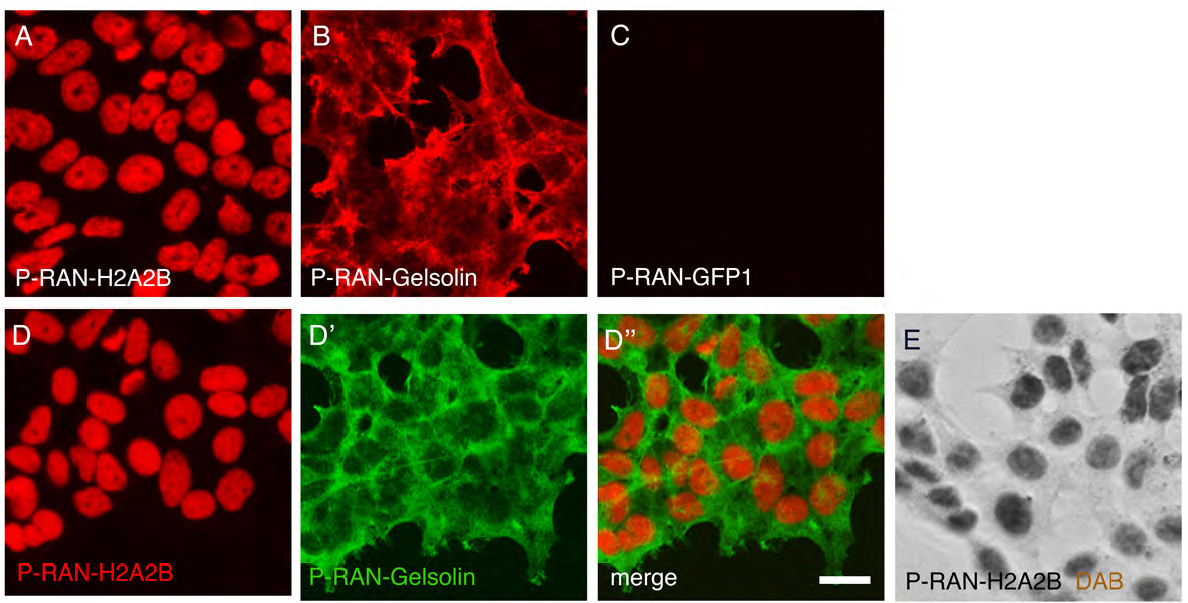
Detection of endogenous proteins with RANbodies. (*A-C*) 293T cells were incubated with P-RAN-H2A2B, P-RAN-Gelsolin or P-RAN-GFP1 as indicated, and stained with Cy3-tyramide for 30 min. Panels were imaged at the same exposure. (*D*) 293T cells were stained with P-RAN-H2A2B/Cy3-tyramide (*D*), stripped with an acidic buffer, and re-stained with P-RAN-Gelsolin/FITC-tyramide (D’). Images are merged in *D*”. (*E*) Untransfected cells incubated with P-RAN-H2A2B and stained with DAB. Bar, 10 *μ*m.

A key advantage of indirect immunohistochemistry is the ability to visualize multiple antigens in the same cells using antibodies from different species. Although this is not possible with RANbodies, we were able to stain with two RANbodies sequentially without cross-reactivity. For this purpose we applied one RANbody, stained with a FITC fluorophore, stripped off the first RANbody with pH 2.0 acidic buffer, and then applied a second RANbody, which we revealed with a Cy3 fluorophore. Fig. 2*D* shows cells doubly-stained with P-RAN-H2A2B and P-RAN-Gelsolin using this protocol.

### Detection of multiple XFPs

We next asked whether RANbodies could be used to amplify signals from XFPs other than GFP and its derivatives (e.g., YFP and Venus). To this end, we generated P-RAN-RFP4, incorporating nanobody LaM-4 described by Fridy et al. (29). P-RAN-RFP4 stained cells transfected with mCherry (a monomeric red fluorescent protein modified from *Discosoma sp* RFP; ref. 30). but did not react with Venus. In contrast, P-RAN-GFP1 stained Venus-transfected but not mCherry-transfected cells (Fig. 3*A* and *B*).

**Fig. 3.**
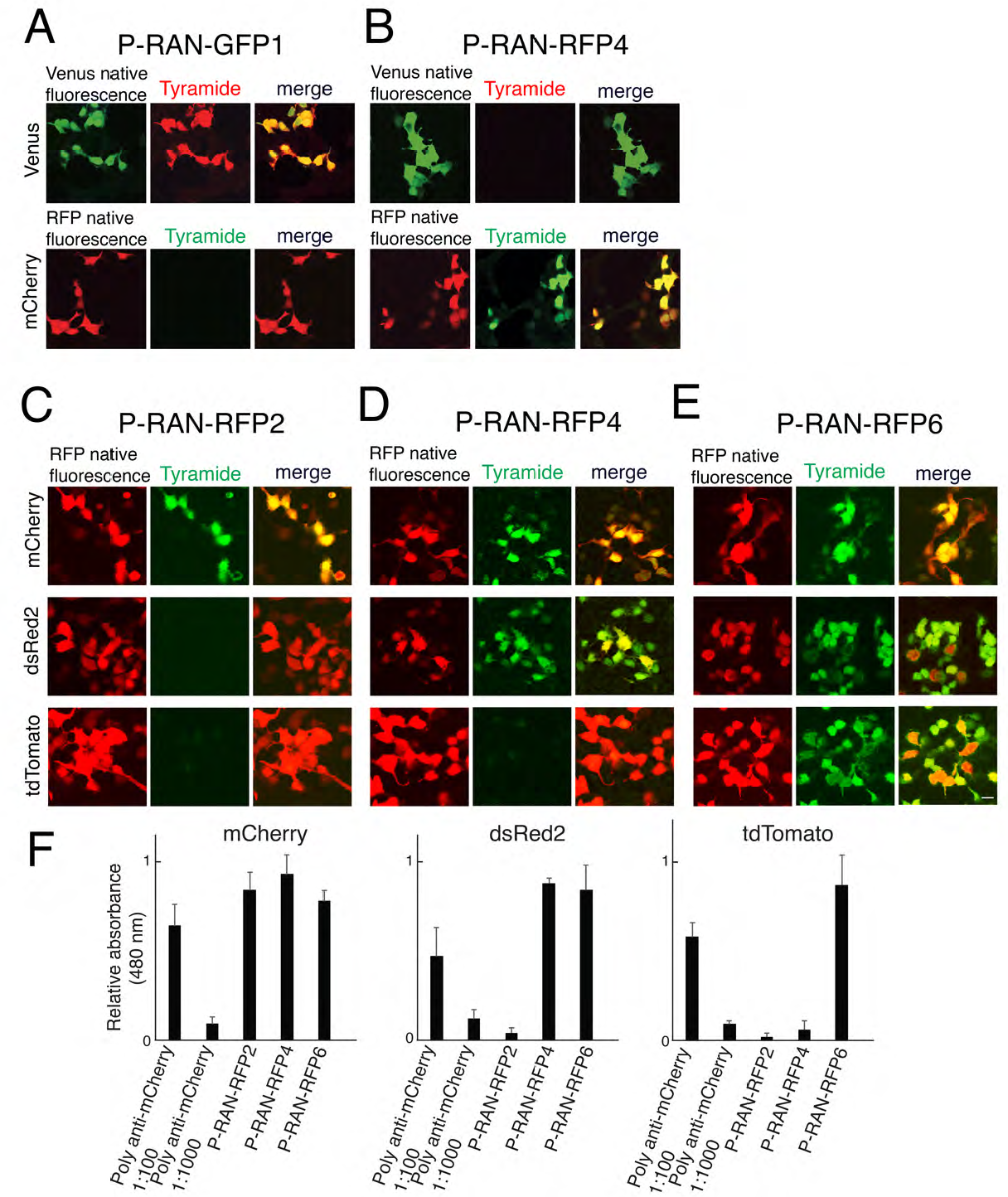
Detection of multiple fluorescent proteins with RANbodies. (*A, B*) 293T cells transfected with Venus or mCherry were incubated with P-RAN-GFP1 (*A*) or P-RAN-RFP4 (*B*), and stained using a Cy3 (red) or FITC (green) tyramide dye. Each RANbody was specific for its cognate antigen. (*C-E*) 293T Cells transfected overexpressing mCherry, dsRed2, or tdTomato (all sea anemone RFP derivatives) were incubated with P-RAN-RFP2 (*C*), P-RAN-RFP4 (*D*), or P-RAN-RFP6 (*E*), and stained with FITC-Tyramide. All P-RAN-RFPs recognize mCherry, but they differ in their ability to detect dsRed2 and tdTomato. Bar, 10 μm. (*F*) Enzymatic detection of cell-bound RANbodies (as in c-e) with a water-soluble substrate provides a second demonstration of the specificity of RFP-RANbodies-P for mCherry, dsRed2, and tdTomato (mean ± SEM, *n=3*). Rabbit anti-RFP polyclonal antibodies react to all the RFP variants.

We also generated two additional RANbodies to RFP, seeking ones that could discriminate amongst red fluorescent proteins. All three, P-RAN-RFP2, -RFP4 and RFP6, derived from LaM-2, -4, and -6, respectively (29), recognized mCherry, but differed in their ability to recognize other RFPs: P-RAN-RFP2 was specific for mCherry, P-RAN-RFP4 recognized both mCherry and dsRed2, a variant of *Discosoma sp* RFP, and P-RAN-RFP6 recognized mCherry, dsRed2 and tdTomato, a tandem dimer of mCherry (Fig. 3*C-F*).

### RANbody-GFP for GRASP amplification

The GFP Reconstitution Across Synaptic Partners (GRASP) method makes use complementation between two fragments of GFP expressed in different cells to label synapses and other sites of intercellular contact (31). Neither of the so-called “split GFP” fragments, sGFP1-10 and sGFP11, is fluorescent on its own, but the reconstituted protein is fluorescent, thus revealing contacts between cells that express different fragments. Because the density of GFP is often low at such sites, the signal is sometimes amplified by use of antibodies that recognize the reconstituted fragment but neither fragment alone (32, 33). However, few such antibodies are available and, for polyclonal antibodies, require affinity purification (33). We therefore asked whether RANbodies could serve as reagents to amplify GRASP signals.

P-RAN-GFP1 recognized both sGFP1-10 and GFP. We therefore generated additional RANbodies to GFP, based on nanobodies LaG-26 and LaG-41 (29) (Table 1). Both P-RAN-GFP26 and P-RAN-GFP41 stained GFP or Venus approximately as well as P-RAN-GFP1 but neither recognized sGFP1-10-linked neurexin (NRXN) or sGFP11-linked neuroligin-1 (NLGN) (Fig. 4*A-C*). We co-transfected sGFP1-10 and sGFP11 into HEK cells to enable intracellular complementation (Fig. 4*D*) and used the enzymatic assay described above to compare the specificity of these GFP RANbodies to that of conventional polyclonal and monoclonal antibodies (Fig. 4*E*). The ability of P-RAN-GFP26 and -GFP41 to discriminate reconstituted GFP from its fragments was superior to that of monoclonal and polyclonal antibodies that have previously been used for this purpose (32, 33). Likewise, P-RAN-GFP26 and -GFP41 can recognize "reconstituted" GFP at cell-cell contacts between sGFP1-10NRXN and sGFP11NLGN transfected cells (Fig. 4*F, G*).

### H, M-RANbodies: RANbodies incorporating highly antigenic reporters

Although P-RANbodies, incorporating HRP, are useful reagents, there are cases in which enzymatic reactions are inconvenient. In addition, tyramide reagents are costly and colorimetric reagents are difficult to incorporate into double- or triple-labeling protocols. We therefore asked whether the HA epitope tag contained in the P-RANbodies (Fig. 1*A*) could be used for immunofluorescent detection. We incubated tdTomato-transfected 293T cells with P-RAN-RFP6, then used an Dylight488-conjugated anti-HA tag antibody to stain the cells. Staining was specific, but dim (Fig. 5*B*). We therefore generated a new set of RANbodies that incorporated highly antigenic reporters called “spaghetti monsters” (18) in place of HRP (Fig. 5*A*). These reporters incorporate 10 HA or MYC epitope tags in a GFP scaffold. As shown in Fig. 5*C, D, M, N, P*, these reporters enabled sensitive detection of RFP and histone (H-RAN-RFP6, H-RAN-H2A2B and M-RAN-H2A2B) either by direct detection using dye-coupled anti-HA antibody or indirect staining with dye-conjugated secondary antibodies. Of the GFP RANbodies tested, H-RAN-GFP26, H-RAN-GFP41 and M-RAN-GFP1 were ineffective because they recognize epitopes in the scaffold (Fig. 5*G*, not shown). However, H-RAN-GFP1 did not recognize the scaffold and was therefore useful for detecting GFP and its derivatives (e.g., Venus) in tissues (Fig. 5*F*).

**Fig. 4.**
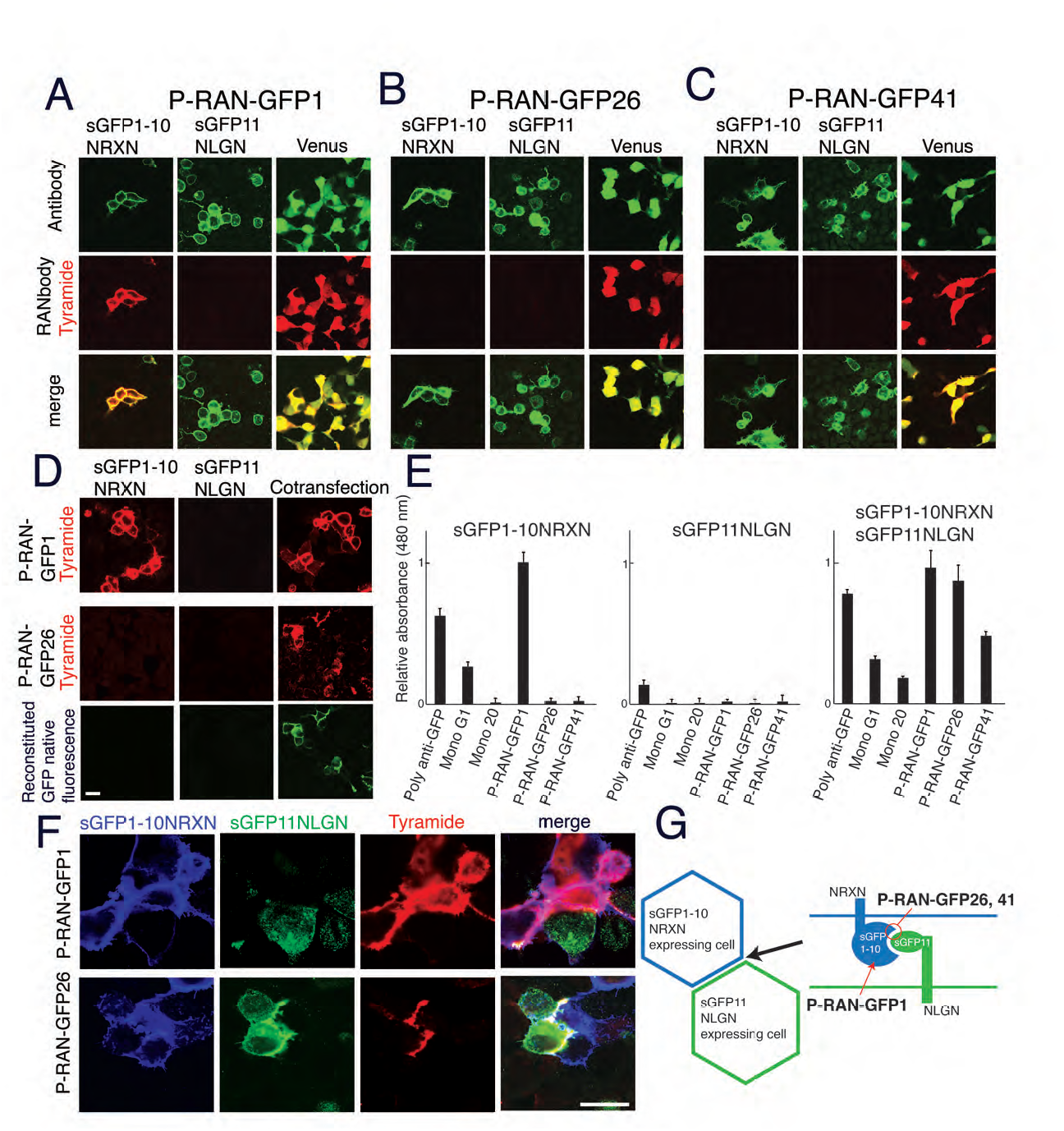
RANbodies-GFP distinguish reconstituted GFP from GFP fragments. (*A-C*) Cells transfected with the 1-10 fragment of GFP fused to neurexin (sGFP1-10NRXN), the short fragment of GFP fused to neuroligin (sGFP11NLGN), or Venus (holo-YFP) were incubated with P-RAN-GFP1 (*A*), P-RAN-GFP26 (*B*), or P-RAN-GFP41 (*C*), and stained with Cy3-tyramide. sGFP1-10NRXN and sGFP11NLGN were detected with anti-NRXN1ß and anti-NLGN1, respectively (top row). All three RAN-GFPs recognize Venus. P-RAN-GFP1 also recognized sGFP1-10, but not sGFP11. P-RAN-GFP26 and -GFP41 do not recognize either fragment. (*D*) sGFP1-10NRXN and sGFP11NLGN plasmids were transfected individually or cotransfected. In the cotransfected cells, reconstituted GFP exhibited green fluorescence, and was recognized by P-RAN-GFP26 as well as P-RAN-GFP1 (Cy3-tyramide). (*E*) Enzymatic assay using a water-soluble HRP substrate confirmed differential reactivity of P-GFP-RANs with sGFP1-10NRXN, sGFP11NLGN, and reconstituted (cotransfected) GFP (mean ± SEM, *n=3*). Mouse monoclonal antibody clone #20 (Mono 20) is selective for reconstituted GFP, whereas monoclonal antibody GFP-G1 and a rabbit anti-GFP polyclonal antibody also detects with sGFP1-10. (*F*) P-RAN-GFP26 was used to detect the reconstituted GFP generated in *trans* at points of contact between cells transfected with sGFP1-10NRXN and sGFP11NLGN and then mixed. P-RAN-GFP1 also stained sGFP1-10NRXN-transfected cells. sGFP1-10NRXN and sGFP11NLGN were detected with chicken anti-GFP (blue) and anti-NLGN1 (green), respectively. (*G*) Schematic of results shown in *F*. Bar in *D*, 10 *μ*m for *A-D*; bar in *F*, 10 *μ*m.

**Fig. 5.**
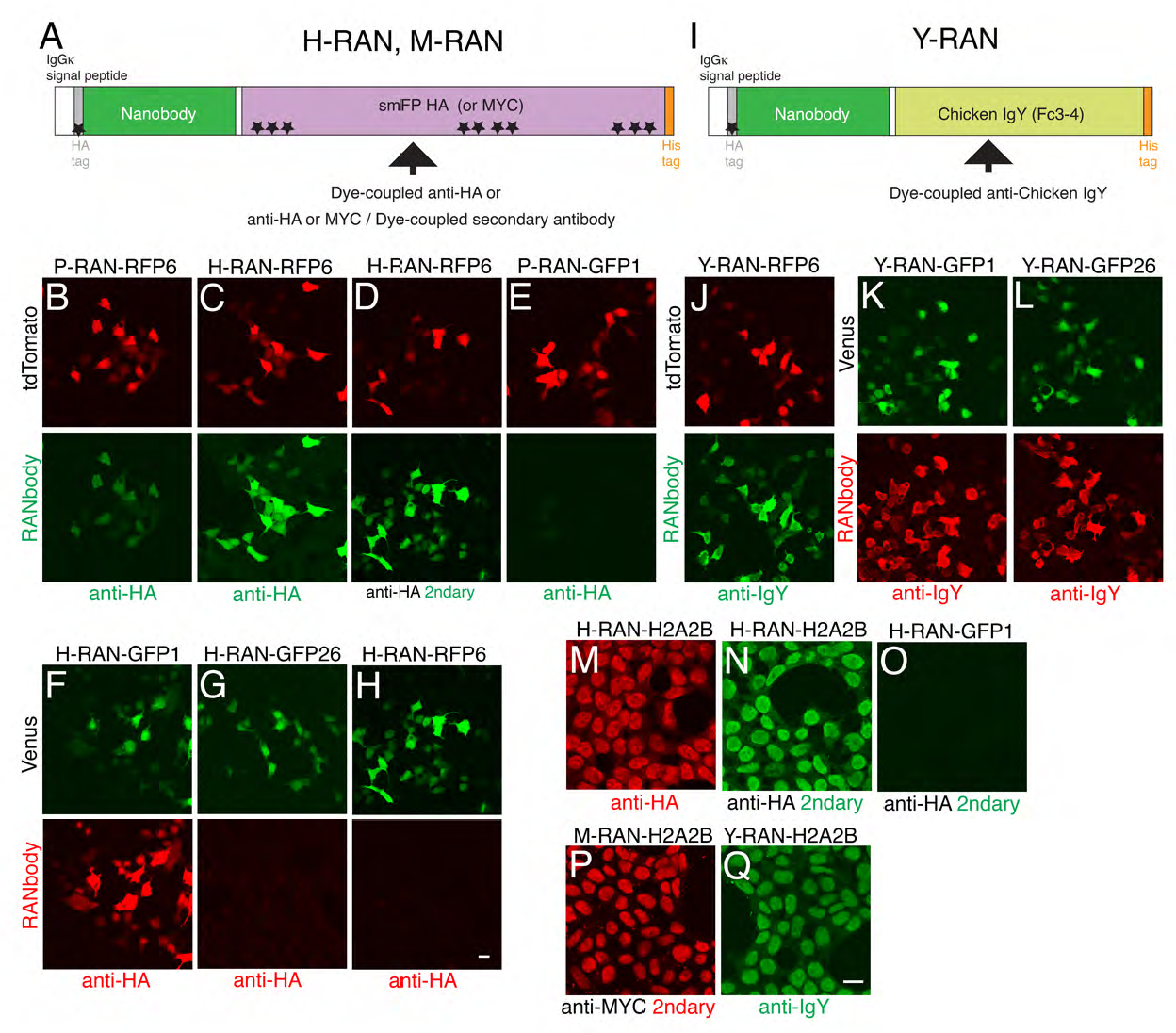
RANbodies incorporating epitope-tagged and antibody-based reporters. (*A*) Structure of H- and M-RANbodies, incorporating 10 HA or MYC epitope tags in a GFP scaffold (spaghetti monster; ref, 18), respectively. These reporters can be stained with antibodies to the HA and MYC epitopes. Note that P-RANbody incorporates a single HA tag. (*B-E*) Detection of tdTomato in 293T cells with P-RAN-RFP6 (with one HA tag) and Dylight488-coupled anti HA (*B*), H-RAN-RFP6 (with 11 HA tags), and Dylight488-coupled mouse monoclonal antibody to HA tag (*C*) or H-RAN-RFP6, rat monoclonal antibody to HA and second antibody (*D*). As a control, tdTomato-transfected cells were incubated with P-RAN-GFP1 and stained with Dylight488-coupled anti-HA (E). All the images were obtained with the same gain. (*F-H*) Venus expressed in 293T cells were incubated with H-RAN-GFP1 (*F*), H-RAN-GFP26 (*G*), or H-RAN-RFP6 (*H*), and stained with Alexa555-coupled anti-HA antibody. H-RAN-GFP1 stained Venus, but H-RAN-GFP26 stained poorly because the nanobody recognized the Spaghetti monster reporter. (*I*) Structure of Y-RANbodies incorporating the Fc3-4 segment of Chicken IgY as reporter. Fc3-4 can be stained with fluorophore-coupled antibodies to chicken IgY. (*J-L*) Detection of tdTomato and Venus in 293T cells with Y series RANbodies, stained with fluorophore-conjugated anti-chicken IgY: Y-RAN-RFP6 (*J*), Y-RAN-GFP1 (*K*) and Y-RAN-GFP26 (*L*). (*M-Q*) Detection of histone in untransfected 293T cells with H-RAN-H2A2B (*M, N*), M-RAN-H2A2B (*P*) and Y-RAN-H2A2B (*Q*), stained as in *C-H*. P-RAN-GFP1 (*O*) serves as a negative control. Bar in H, 10 *μ*m for *B-H*; bar in *Q*, 10 *μ*m for *M-Q*.

### Y-RANbody: RANbodies incorporating a chicken Fc fragment

In many cases, mouse or rat monoclonal and rabbit polyclonal antibodies are the only available reagents for detecting antigens in tissue. Detection of additional antigens with conventional second antibodies therefore requires use of antibodies derived from other species. To expand the possibilities for multiple-labeling, we used an Fc fragment of chicken IgY as a reporter (Fig. 5*I*). This Fc3-4 fragment contains two immunoglobulin domains, which can be detectable with readily-available fluorophore-coupled secondary antibodies to chicken IgY (Fig. 5*J, K, L, Q*). Indeed, RANbodies incorporating this reporter, Y-RAN-RFP6, Y-RAN-GFP1, Y-RAN-GFP26 and Y-RAN-H2A2B were approximately as sensitive as RANbodies incorporating spaghetti monsters as reporters (Fig. 5). Moreover, unlike spaghetti monster-derived H-RAN-GFP26, the IgY-derived Y-RAN-GFP26 was able to detect GFP (Fig. 5*L*).

### Detection of antigens with RANbodies in tissue sections

Finally, we assessed the ability of RANbodies to stain tissue sections. We used mouse retina because we have characterized several transgenic lines in which retinal neurons express XFPs. P-RAN-GFP1 recognized GFP derivatives in each of two such lines tested. In TYW3, subsets of retinal ganglion cells (RGCs) express YFP cytoplasmically, revealing somata and dendrites of several distinct RGC types (34, 35). In Sdk1::CreGFP mice (M.Y. and J.R.S., in preparation), CreGFP is localized to the nuclei of Sdk1-expressing cells, some of which have been characterized previously (36). Sections were treated with H_2_O_2_ to inactivate endogenous peroxidase-like activities, then incubated with P-RAN-GFP1 and reacted with the tyramide substrate. In both lines, staining was more intense with RANbody than with conventional anti-GFP antibodies and fluorophore-conjugated second antibodies (Fig. 6*A-H*). In our hands, Cy3-Tyramide (red color) resulted in crisper staining with lower background than FITC-tyramide (Fig. 6*C, D, G, H*) and sensitive staining than chromogenic DAB staining (not shown). Moreover, fine dendrites in the inner plexiform layer were more clearly visualized with P-RAN-GFP1 than by indirect immunofluorescence (Fig. 6*L*). Similarly, P-RAN-RFP6 stained tdTomato-positive cells in double transgenics (Parvalbumin-Cre mated to Ai14, a tdTomato reporter line; refs, 37 and 38). Again, RANbody staining was more intense and better defined than indirect immunofluoresences using anti-tdTomato (Fig. 6*I-K*). The staining is compatible with immunohistochemistry using monoclonal or polyclonal antibodies with anti-mouse or rabbit secondary antibodies (see, for example, Fig. 6*M*).

**Fig. 6.**
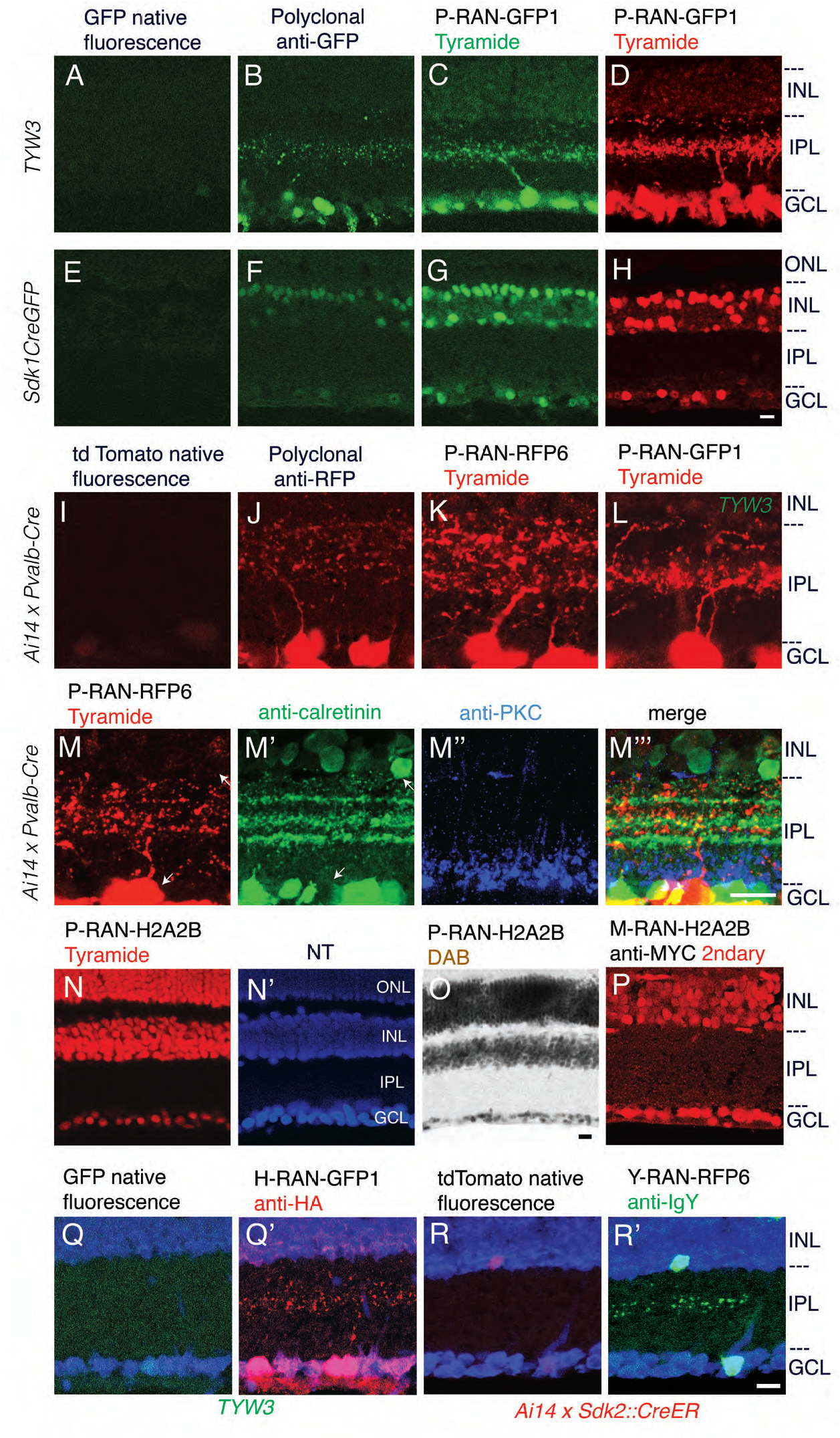
Detection of antigens in tissue sections by RANbodies. (*A-D*) Sections of retina from a TYW3 mouse, in which YFP is expressed in subsets of retinal ganglion cells with dendrites in the central sublaminae of the inner plexiform layer (IPL). A, Native fluorescence; *B*, chicken anti-GFP; *C*, P-RAN-GFP1/FITC-tyramide; *D*, P-RAN-GFP1/ Cy3-tyramide. All the images were obtained with the same gain. ONL, outer nuclear layer; INL, inner nuclear layer; GCL, ganglion cell layer. (*E-H*) Sections of retina from a Sdk1::CreGFP mouse, in which CreGFP is localized to nuclei of subsets of cells in the GCL and INL. Staining as in *A-D*. (*I-K*) Sections of retina from a double transgenic (cre-dependent tdTomato;Parvalbumin-cre) in which subsets of retinal ganglion cells express tdTomato. Sections were stained with anti-RFP or P-RAN-RFP6/Cy3-tyramide. The RANbody stained dendrites more distinctly than the antibody. (*L*) TYW3 retina stained with P-RAN-GFP1 as in A-D, showing similarly distinct staining of dendrites. (*M*) Section of retina from a cre recombinase-dependent tdTomato;Parvalbumin-cre mouse stained with P-RAN-RFP6/Cy3-tyramide, rabbit anti-calretinin, mouse antiprotein kinase C (PKC), and anti-rabbit and mouse secondary antibodies. RANbodies can be used together with conventional antibodies from multiple species. (*N,O*) Sections of wildtype mouse retina incubated with P-RAN-H2A2B and stained with Cy3-tyramide (*N*) plus neurotrace (*N*’) or DAB (*O*). The RAN-H2A2B stained all nuclei. (*P*) Section of wildtype mouse retina incubated with M-RAN-H2A2B, stained with anti-MYC. All nuclei are stained. (*Q,Q*’) Section of adult retina from TYW3 mouse stained with H-RAN-GFP1 /Alexa 555-coupled mouse monoclonal antibody to HA tag (2-2.2.14). (*R,R*’) Section of adult retina from Ai14;Sdk2::CreER stained with Y-RAN-RFP6 and Alexa488-conjugated anti-chicken IgY secondary antibodies. In both *Q* and *R*, neuronal processes in the inner plexiform layer (IPL) are clearly visible using RANbodies (*Q*’, *R*’) whereas native fluorescence from reporters (GFP and tdTomato) is inadequate to reveal these processes (*Q, R*). Bar in *H*, 10 μm for A-H; bar in *M*"' 10 μm for *I-M*"'; bar in O, 10 *μ*m for *N-O*; bar in *R*', 10 *μ*m for *P-R*'.

We also stained mouse retina with P-RAN-H2A2B and observed strong, specific staining of nuclei (Fig. 6*N, O*). Thus, RANbodies can be used to detect endogenous antigens in tissue.

Finally, we confirmed that RANbodies incorportating HA-based spaghetti monsters, MYC-based spaghetti monster, and a chick IgY Fc fragment (H, M, and Y-RANbodies, respectively) could all be used to reveal antigens in tissue sections. Examples are shown in Fig. 6*P-R*.

## DISCUSSION

This paper describes RANbodies, versatile reagents generated by fusing a reporter to a nanobody. RANbodies can be used to detect antigens in cells and tissues. They are sensitive, because of the amplification provided by the reporters (enzymatic for HRP and incorporation of multiple epitopes in the spaghetti-monsters); specific, because of the “monoclonal” nature of nanobodies; inexpensive, because they can be produced by standard methods in most laboratories; and “eternal” because they can be regenerated based on information contained in a simple digital sequence file. We show that they can be used to detect a variety of antigens in both cultured cells and tissue sections. In addition to detecting endogenous antigens (e.g., histones and gelsolin), they can be used to amplify the endogenous fluorescence of GFP, RFP, tdTomato and mCherry, decreasing reliance on expensive commercial anti-XFP antibodies.

We generated RANbodies incorporating four reporters, allowing visualization by using colorimetric and fluorescent enzymatic amplification (P series, HRP), commercially available anti-epitope tag antibodies (H and M series, HA- and MYC-tagged spaghetti monsters) and second antibodies (Y series, chicken IgY). Each has advantages. The amplification provided by the enzymatic activity of HRP renders the P series many-fold more sensitive than standard indirect fluorescence. Moreover, no secondary antibody is required for detection. On the other hand, enzymatic reaction on tissue is sometimes cumbersome and requires careful monitoring of incubation time. Moreover, the tyramide reagents used for fluorescent detection are expensive, and the inexpensive colorimetric substrates are incompatible with most multiple labeling protocols. Conversely, H-, M-, and Y-RANbodies are simpler to use and well suited for multiple labeling but, although they are approximately as sensitive as standard indirect immunofluorescence, they are less sensitive than P-RANbodies. The Y-RANbodies have the additional of enabling use of anti-chicken second antibodies, which can be combined with more widely used anti-rodent and –rabbit secondary antibodies. For example, to our knowledge, the best currently available antibodies to tdTomato, are generated in rabbits, and are therefore difficult to combine with broadly available rabbit antibodies. Y-RAN-RFPs provides a useful alternative, increasing the range of antibodies that can be used in combination with RFP detection.

The advantages of nanobodies have been broadly appreciated: a search of Pubmed with the term “nanobody OR nanobodies” retrieves 617 items in 2015-2017 alone. Thus, new nanobodies are being generated at a rapid rate. As nanobodies to additional antigens become available, the utility of the RANbody method is likely to increase.

## MATERIALS AND METHODS

### Construction of RANbodies

DNA sequences including a signal sequence from the human immunoglobulin kappa chain, a hemagglutinin (HA) tag, codon-optimized HRP, and a His tag were assembled in a backbone of pCMV-N1-EGFP (Clontech, Mountain View, CA) together with sequences encoding nanobodies which had been synthesized as gBlock gene fragments (IDT, Coralville, IA) using the Gibson Assembly HiFi 1-Step kit (SGI-DNA, La Jolla, CA).

In some cases, the synthesized nanobody sequences were inserted between the HA tag and HRP sequence after amplifying the vector sequence using primers GACTATCCATATGATGTTCCAGATTATGCT (reverse strand at HA tag) and GGAACACAGGTGACGGTGAGCTCTAGCGGT (forward strand at the 5' end of HRP) and high fidelity Q5 DNA polymerase (New England Biolabs, Ipswich, MA).

In other cases, reporter sequences were inserted between the nanobody and His tag sequence after amplifying the vector sequence using primers ACCGCTACCGCTAGAGCTCACCGT (reverse strand at nanobody-reporter junction) and TTGCATCATCATCATCATCATCATTGA (forward strand at His tag) and high fidelity Q5 DNA polymerase. Insertions were as follows. P-RANbody: ---SSSGSG<u>MQLTPTFYDNSC---GQIRRNCRVVNSNSD</u>LHHHHHHH (underlined sequence corresponds to vHRP, see Fig. 1*B* and Table 1 for details) H-RANbody: ---SSSGSG<u>MYPYDVPDYAG---- AGYPYDVPDYA</u>LHHHHHHH (underlined sequence from mFP-HA, obtained from Addgene, GenBank KR107022) M-RANbody: ---SSSGSG<u>MEQKLISEEDLAEQ---- LAEQKLISEEDL</u>LHHHHHHH (underlined sequence from mFP-Myc, obtained from Addgene; GenBank KR107023) Y-RANbody: ---SSSGSG<u>PGELYISLDAKLRCLVVNLPS----FSQRTLQKQAGK</u>LHHHHHHH (underlined squence from chicken IgY-Fc3-4 fragment, GenBank: X07174)

Sequences for nanobodies described in this report are presented in Table 1. RANbody constructs are available from Addgene.

### Other plasmids

mGRASP constructs (sGFP1-10NRXN, sGFP11NLGN, sGFP1-10NLGN, sGFP11NRXN; available from Addgene) were described previously (33). Other XFPs used in this report are as follows: Venus, an enhanced yellow fluorescent protein modified from *Aequorea victoria* GFP (26); Cerulean, an enhanced blue fluorescent protein derived from *Aequorea victoria* GFP (39); dsRed2 (Clontech), a derivative of potentially tetrameric red fluorescent protein modified from *Discosoma sp* red fluorescent protein (40); mCherry, a monomeric red fluorescent protein modified from *Discosoma sp* red fluorescent protein (30); tdTomato, a tandem dimer red fluorescent protein generated from *Discosoma sp* red fluorescent protein (41); mKate2 (Evrogen), a far-red mutant fluorescent protein derived from *Entacmaea quadricolo* fluoresenct protein (Shcherbo et al., 2009).

### Cell lines and transfection

Human embryonic kidney (HEK) 293T were purchased from the American Type Culture Collection (ATCC; Manassas, VA). Since HEK-293T cells are often contaminated with HeLa cells (International Cell Line Authentication Committee; http://iclac.org/databases/cross-contaminations/), we confirmed that the cells were G418 resistant, a characteristic of 293T but not HeLa cells. Cells were cultured in Dulbecco Modified Eagle Medium (DMEM) supplemented with 10% (v/v) fetal calf serum and penicillin/streptomycin plus Normocin and G418 (Invivogen, San Diego, CA) (DMEM10). In some cases, culture dishes were coated with Attachment Factor Protein (ThermoFisher/Invitrogen).

For transient expression, 293T cells were transfected with DMRIE-C (ThermoFisher/Invitrogen) in OptiMEM (ThermoFisher/Invitrogen), and incubated in DMEM10 for 3 days. The cells were trypsinized, suspended in DMEM10 supplemented with 10 *μ*g/ml deoxyribonuclease I (DN25, Sigma), and cultured in the same medium (with DNAse) on glass coverslips (Bellco) coated with Attachment Factor Protein. For studies in which cells transfected with different constructs were to be mixed, the transfected cells were treated with deoxyribonuclease I before mixing populations to avoid transfer of extracellular plasmid transfers.

To generate 293TA lines stably expressing split GFP constructs, the sequences were amplified with Q5 DNA polymerase (New England Biolabs, Ipswich, MA), digested with restriction enzymes, and cloned into a piggyBac transposon vector pXL-CAG-Zeocin-3xF2A (Yamagata and Sanes, in prep; Martell et al., 2016; Goodman et al., 2016). Cells were transfected with an appropriate construct together with a piggyBac transposase vector pCAG-mPBorf (33) using DMRIE-C, trypsinized after 2 days, replated into larger plates, and selected with 1 mg/mL Zeocin (Invivogen, San Diego, CA) for 1-2weeks. Surviving colonies were transferred to new plates and screened with antibodies against cognate molecules with high and homogeneous expression. These stably-transfected cells were trypsinized, and plated on the attachment factor protein-coated glass coverslips. In some experiments, cells were infected with VSV-G-pseudotyped lentiviral vectors that overexpress sGFP1-10NRXN or sGFP11NLGN under CMV promoters.

### Antibodies

Chicken antibodies to GFP, mouse monoclonal antibody to GFP, rabbit anti-mCherry and rabbit anti- lacZ were generated in our laboratory (33, 42, 43). The monoclonal antibody to GFP, GFP-G1, is available from Developmental Studies Hybridoma Bank, Iowa City, IA). Other primary antibodies were as follows: Mouse monoclonal anti-GFP *#*20 (Sigma-Aldrich, St. Louis, MO), rat anti-HA tag (clone 3F10; Roche, Indianapolis, IN), rabbit anti-calretinin (Millipore, Bedford, MA), mouse monoclonal anti-protein kinase C alpha (clone*#* MC5; Sigma), goat anti-human CD4 (RD systems, Minneapolis, MN), mouse monoclonal anti-neuroligin-1 (clone*#* N97A/31, NeuroMab, Davis, CA) and neurexin-1β (clone*#* N170A-1, NeuroMab, Davis, CA). Alexa Fluor 555- and DyLight 488 conjugated HA Tag Monoclonal Antibody (2-2.2.14) were from Invitrogen. Mouse monoclonal anti-Myc antibody (9E10) was from DSHB, Iowa City, IA. Fluorophore-conjugated secondary antibodies were obtained from Jackson ImmunoResearch (West Grove, PA). Neurotrace 640 was from ThermoFisher/Invitrogen.

### Mice

The Ai14 and parvalbumin-cre line (Pvalb^tm1(cre)Arbr^) lines were obtained from Jackson Laboratories. Ai14 expresses tdTomato in a Cre-dependent manner (Rosa26-CAG-lox-stop-lox-tdTomato) (38). In Pvalb-cre, an IRES-Cre cassette was inserted downstream of the coding sequence in the endogenous parvalbumin locus (37). TYW3 mice, Sdk1::CreGFP, Sdk2::CreER-T2 mice were generated in our laboratory. In TYW3, lox-flanked EYFP is expressed under control of regulatory elements from the Thy1 gene (34). In Sdk1::CreGFP, a Cre-GFP was inserted at the translational start site of the *Sdk1* gene using CRISPR/Cas9 genome engineering (M.Y. and J.R.S., in preparation). Sdk2::CreER-T2 mouse line was previously described (36). CD1 mice were purchased from Jackson Laboratories. Animals were used in accordance with NIH guidelines and protocols approved by Institutional Animal Use and Care Committee at Harvard University.

### Production of RANbodies

To produce RANbody proteins, 293T cells cultured on tissue culture dishes precoated with Attachment Factor Protein (ThermoFisher/Invitrogen) were transfected with each RANbody plasmid DNA using a calcium phosphate precipitation method followed by 15%(w/v) glycerol shock in serum-free DMEM. After switching to DMEM10 or Opti-MEMI (ThermoFisher/Invitrogen), the cells were incubated for 3 days. The medium, which contained secreted RANbody, was harvested, filtered through 0.45*μ*m pore cellulose acetate membranes, and applied to cobalt Talon resin columns (Clontech), rinsed, and eluted according to the manufacturer’s protocol. The eluate was concentrated using Pierce 9K concentrators (Thermo), and then substituted with several cycles of PBS. RANbodies were stored at 4C° with 0.01% (v/v) ProClin 150 (Sigma-Aldrich) as a preservative, or aliquoted and frozen at −20°C.

For usage, RANbodies were generally not purified to homogeneity, so concentration cannot be determined accurately, but we estimate it to be 0.1-1*μ*g/ml in most cases. In practice, it is possible to use cultured medium harvested after plasmid transfection without further purification.

### Staining and imaging

Cultured cells were fixed with 4%(w/v) paraformaldehyde/PBS at 4°C for 15 min, treated with 0.1%(w/v) Triton X100/ PBS supplemented with 0.3%(w/v) H_2_O_2_ at 4°C for 15 min, rinsed with PBS, and blocked with 5%(w/v) skim milk in PBS at room temperature for 30 min. After removing the blocking solution, the cells were incubated with RANbody, diluted in DMEM10 overnight at 4°C. After rinsing with PBS, cells that had been incubated with P-RANbodies were stained with tyramide dye (TSA Plus kit, Perkin Elmer, Waltham, MA), rinsed overnight with PBS, and mounted in Fluoro-Gel (Electron Microsopy Sciences). Staining with 3,3'diaminobenzidine (DAB) was performed using DAB peroxidase substrate kit SK-4100 (Vector Laboratories, Burlingame, CA). H-, M-, and Y-RANbodies were stained with antibodies and second antibodies listed above.

Immunostaining was performed after staining with a tyramide dye using appropriate primary secondary antibodies. To carry out successive staining with two different RANbodies, the first RANbody was removed by treating cells with 0.1M Glycine, pH2.0 for 30 min at room temperature after development of the first tyramide reaction, then incubated with the second RANbody followed by the second color tyramide reaction.

For staining retina, mice were anesthetized and perfused with Hanks’ balanced salt solution. Retinas were then dissected, fixed with 4%(w/v) paraformaldehyde/PBS for 10 hours, sunk in 15%(w/v) and 30%(w/v) sucrose, frozen in the Tissue-Tek O.C.T. compound (Electron Microscopy Sciences, Hatfield, PA), cryosectioned, collected on Superfrost Plus slides (VWR Scientific, Franklin, MA), and dried at room temperature. For P-RANbodies, sections were pretreated with 0.1% (w/v) Triton X100/PBS plus 0.3% (w/v) H_2_O_2_ at room temperature for 3 hours, rinsed with PBS, and blocked with 5% (w/v) skim milk /PBS at room temperature for 30 min, and stained as described above. Imaging was done with a Zeiss Meta510 confocal microscope. For H-, M-, and Y-RANbodies, sections were stained with antibodies and second antibodies listed above. Images were processed with Adobe Photoshop, and Image-J (Version 1.47d, Fiji).

### Enzyme assays

To compare the activity of HRP variants, transfected cells were lysed with 0.1% (w/v) TritonX-100/PBS. The lysate was incubated with o-phenylenediamine/H_2_O_2_, stopped with 8M sulfuric acid, and quantified using a spectrophotometer at 480nm.

To quantify HA-tagged molecules in the lysate, the lysate was coated onto plastic wells, and colorimetric enzyme-linked immunosorbent assay (ELISA) was carried out using anti-HA tag antibody and alkaline-phosphatase-conjugated anti-rat secondary antibodies. The wells were developed with *p*-nitrophenyl phosphate/50mM MgCl_2_/50mM diethanolamine, pH 9.5; and measured at 405 nm.

To detect cell-bound RANbody, paraformaldehyde-fixed cells were treated with 0.3% H_2_O_2_/PBS for 30 min at room temperature, blocked with 5% (w/v) skim milk/PBS for 30 min, incubated with RANbodies overnight, rinsed PBS and developed with *o*-phenylenediamine/H_2_O_2_ as above. Mouse and rabbit antibodies were assayed similarly, except that peroxidase-conjugated secondary antibodies (Jackson ImmunoResearch, 1:1000 dilution in 0.5% BSA/PBS) were added for 2 hr following the first incubation.

## ACKNOWLEDGEMENTS

This work was funded by NIH R37 NS029169.

## CONTRIBUTIONS

MY performed all the experiments. MY and JRS designed the study and wrote the manuscript.

## SUPPLEMENTARY FIGURE LEGENDS

**Fig. S1.**
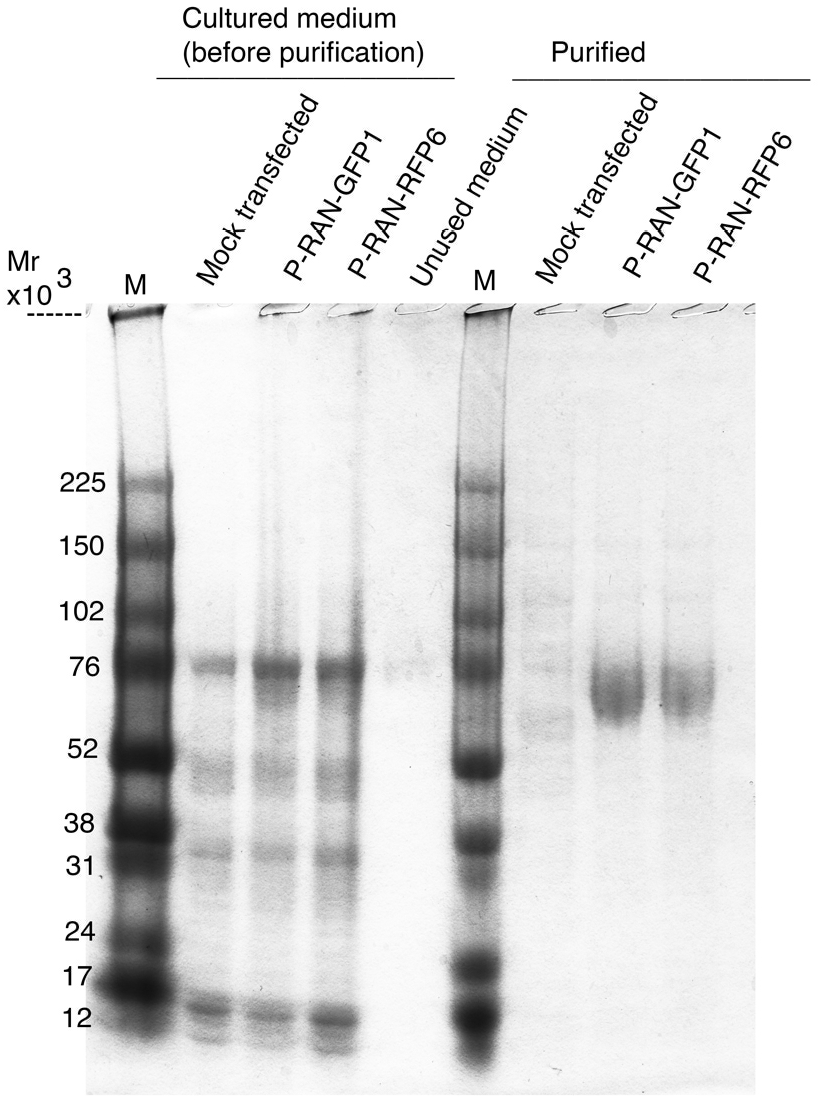
SDS-PAGE analysis of RANbodies purified from transfected 293T cultured in serum-free medium. Each lane corresponds to ~1 ml of medium (left) or RANbody purified from ~1 ml of medium. M, molecular weight markers. RANbody Mr is ~65kDa. The diffuse nature of the bands likely reflects heterogeneous glycosylation.

**Fig. S2.**
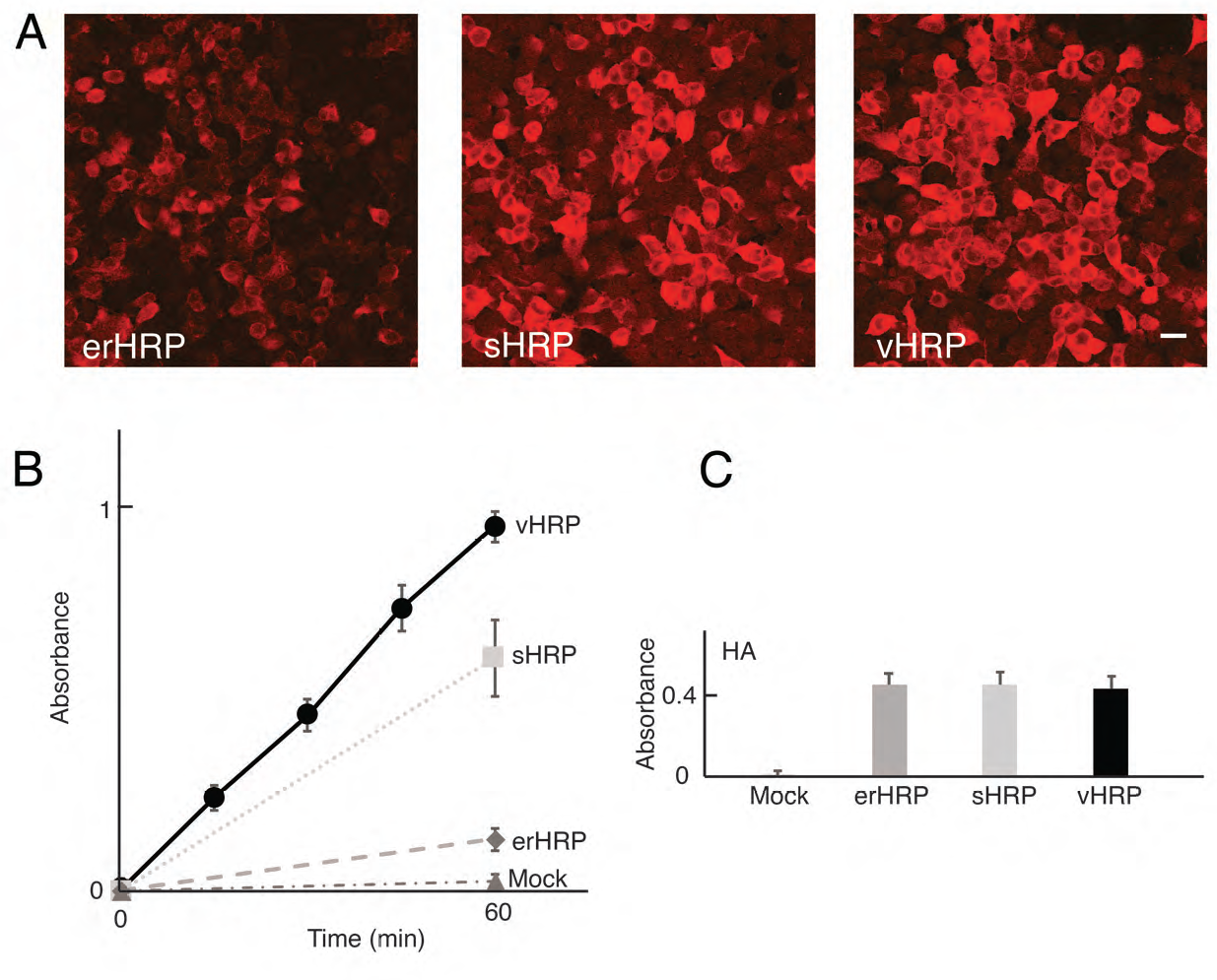
(*A*) 293T cells transfected with HRP variants and reacted with Cy3 tyramide. erHRP: codon-optimized HRP with an endoplasmic reticulum retention signal; sHRP: erHRP with an N175S mutation; vHRP: sHRP with 5 additional mutations (see Fig. 1*B*). All were reacted and photographed under identical conditions. Bar, 10 μm. (*B*) Peroxidase activity of the three HRP variants assayed in lysates from the transfected cells (mean ± SEM, *n=3*). (*C*) Relative levels of HRP variants in lysates, assayed with anti-HA and an alkaline-phosphatase activity (mean ± SEM, *n=3*).

**Fig. S3.**
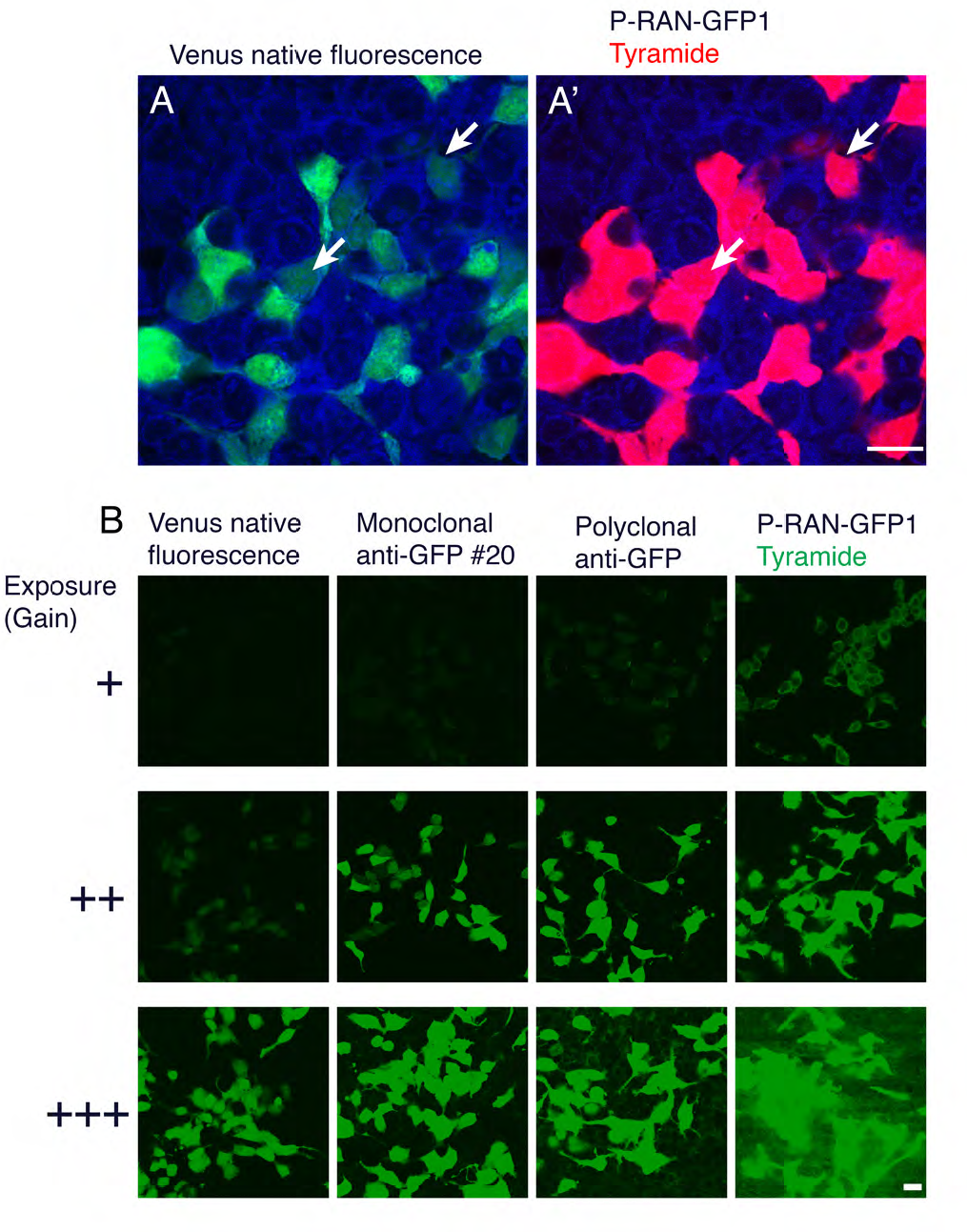
Sensitivity of P-RAN-GFP1 and anti-GFP antibodies. (*A*) 293T cells transfected with Venus were stained with P-RAN-GFP1 and Cy3-tyramide (*A*, native fluorescence; *A*’, Cy3). Some cells with barely detectable native fluorescence were readily detectable with RANbody staining (arrows). The cells were counterstained with NeuroTrace 640 (blue). Bar, 10 *μ*m. (*B*) Cells transfected with Venus were stained mouse anti-GFP (clone *#*20) plus Alexa488-coupled donkey anti-mouse IgG, chicken anti-GFP plus Alexa488-coupled donkey anti-chicken IgY(1:1000 dilution), or P-RAN-GFP1 (1:10 dilution) plus FITC-tyramide (developed for 3 hrs). Cultures were then imaged at three exposures (gains), with gains identical for each row (+, ++, and +++). Only RANbody staining is clearly detectable at the lowest gain and native fluorescence is easily detectable only at the highest gain. Bar, 10 *μ*m.

**Supplementary Table 1.**
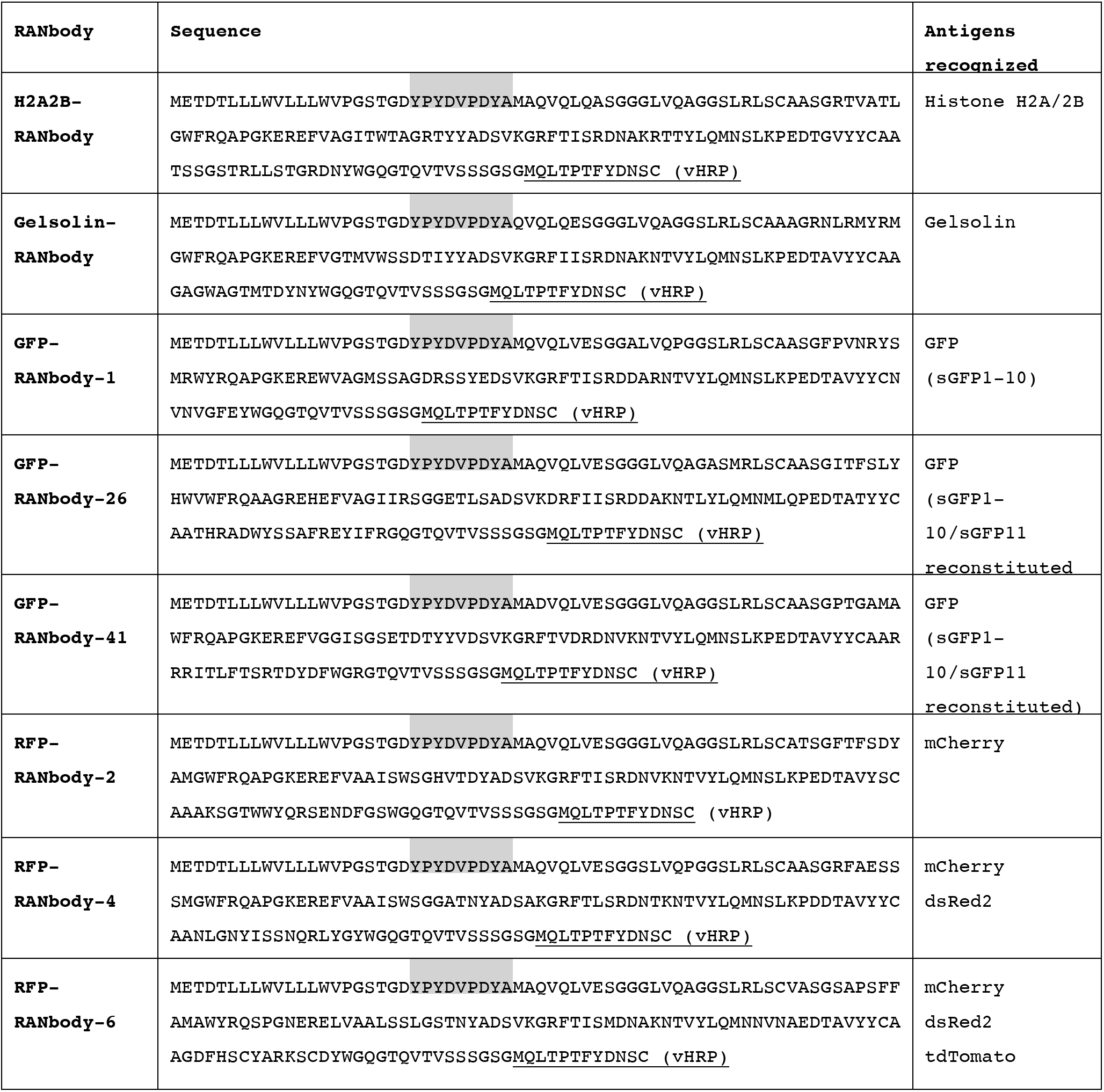
Amino acid sequences of RANbodies generated for this study. The Table shows the sequence of RANbodies, from the amino terminal to the first 11 amino acids of vHRP (underlined). Shaded amino acids (YPYDVPDYA) correspond to a HA-tag after the common signal sequence (see Figure 1*B*). Nanobody sequences are between the shaded and underlined blocks. Sequences were from previous reports (Histone H2A/H2B heterodimer, Jullien et al., 2016; Gelsolin, Van den Abbeele et al., 2010; GFP-nanobody-1, Kubala et al., 2010; other FP nanobodies, Fridy et al., 2014).

